# miRNA binding pressure channels evolution of SARS-CoV-2 genomes

**DOI:** 10.1101/2023.03.31.535057

**Authors:** A. Zhiyanov, M. Shkurnikov, A. Nersisyan, H. Cai, A. Baranova, A. Tonevitsky

## Abstract

In somatic cells, microRNAs (miRNAs) bind to the genomes of RNA viruses and influence their translation and replication. Here we demonstrate that a significant number of miRNA binding sites locate in the NSP4 region of the SARS-CoV-2 genome, and the intestinal human miRNAs exert evolutionary pressure on this region. Notably, in infected cells, NSP4 promotes the formation of double-membrane vesicles, which serve as the scaffolds for replication-transcriptional complexes and protect viral RNA from intracellular destruction. In three years of selection, the loss of many miRNA binding sites, in particular, those within the NSP4, has shaped the SARS-CoV-2 genomes to promote the descendants of the BA.2 variants as the dominant strains and define current momentum of the pandemics.

## 1 Introduction

The (+)RNA genome of the pandemic SARS-CoV-2 virus is about 30,000 nucleotides long [1]. Since December 2019, when the original strain (Wuhan-Hu-1) has emerged, the SARS-CoV-2 acquired more than 10,000 recorded nucleotide mutations [2], with an estimated mutation rate between 0.0004 and 0.002 mutations per nucleotide per year [3]. The current release of the GISAID database contains more than 14 million SARS-CoV-2 sequences [4] and allows the tracking of the evolution of the virus with unprecedented precision. As widely discussed, SARS-CoV-2 is being shaped by the pressure exerted by neutralizing antibodies and applied upon the receptor-binding domain of the Spike (S) protein [5]. It should be, however, noted that antibodies are far from being the only adversarial factor encountered by the virus upon infection. In particular, the binding of host miRNA to other (+)RNA viruses provides a substantial evolutionary pressure [6] as these molecules suppress both the translation and the replication of virus along with altering some aspects of their pathogenesis [7–9].

Several works have predicted interactions of the SARS-CoV-2 single-stranded (+)RNA virus with multiple human miRNAs [10–13]. A majority of studies have not, however, taken into account the ability of the SARS-CoV-2 virus to evolve, and, accordingly, to selectively alter its miRNA binding regions, especially these enriched in binding seeds. Only a few published works utilized the sequences from various Variants of Concern (VOCs) [14].

Human genomes encode many miRNAs, a majority of which display differential expression in human tissues. It is obvious that only miRNA species expressed in replication-permissive host cells will be relevant to SARS-CoV-2 evolution. In humans, the SARS-CoV-2 virus predominantly replicates in the type 2 alveocytes of the lung. In addition, significant clinical data indicate the presence of the virus in the intestinal epithelium, which expresses both ACE2 and TMPRSS2 [15]. Moreover, intestinal viral persistence is commonly detected. It is now recognized that long-term residence of SARS-CoV-2 in the human gut supports low-intensity inflammation, which contributes to the complications of COVID-19, collectively known as post-COVID or long COVID [16].

For three years of the pandemic, the SARS-CoV-2 virus has significantly mutated. Due to the degeneracy of the genetic code, many mutations have not led to a change in the sequence of viral proteins and, therefore, are classified as silent [17]. Nevertheless, silent nucleotide changes have managed to get fixed in the virus genome. In evolution, the fixation of the mutation may take place either by chance, or under the influence of the selective forces acting upon RNA itself rather than on the encoded proteins. The binding of the cellular miRNAs to the viral genome may be capable of exerting such pressure. Here we tested the hypothesis that the evolutionary pressure exerted by binding of miRNA species expressed in human lung and intestinal tissue has, in fact, shaped up the genome of the current variants of SARS-CoV-2 virus dominating the globe.

## 2 Materials and methods

### 2.1 SARS-CoV-2 variants

A total of 14,472,180 SARS-CoV-2 RNA sequences were downloaded from the GISAID database [18] on January 7^th^ 2023 and annotated with Pangolin [19] (program’s version 4.2, Pangolin’s data version 1.17). Among 28,420 variants associated with the City of Berlin (Federal Republic of Germany), we selected 3,755 ones satisfying the following conditions: viral RNA does not contain unrecognized nucleotides; the difference between viral RNA length and the “Wuhan” reference variant (GISAID: EPI ISL 402125) is less than 5%. All selected RNA sequences were collected between the 14^th^ January 2021 and 12^th^ December 2022.

All sequences were divided into two groups, “Ancestral variants” and “Omicrons”, according to their GISAID descriptions. “Ancestral variants” included variants of “Alpha”, “Beta”, “Delta”, and others (Table 1 of Supplementary materials) and collected before the 30^th^January 2022, while all “Omicrons” species were sequenced after 1^st^December 2021 (Figure 1).

**Table 1:** Statistically significant differences in the weighted number of colon miRNA binding motifs separated by SARS-CoV-2 RNA coding regions. All weighted numbers are averaged by variants of the groups and normalized by “Wuhan” RNA length. “Diff” columns contain differences between the averaged values of the corresponding groups. “FDR” columns contain adjusted p-values of Mann-Whitney U tests comparing the weighted numbers of virus variants of the corresponding groups.

| Region | Length | Ancestral variants | Omicron BA.2-like |  | Other Omicrons |  |
| --- | --- | --- | --- | --- | --- | --- |
|  |  |  | Diff | FDR | Diff | FDR |
| NSP4 | 1500 | 962 | 464 | < 0.0001 | 1 | 0.1127 |
| Spike | 3822 | 709 | 21 | < 0.0001 | 20 | < 0.0001 |
| NSP15 | 1038 | 376 | 21 | < 0.0001 | 11 | < 0.0001 |
| NSP9 | 339 | 1518 | 6 | < 0.0001 | 7 | < 0.0001 |
| NSP3 | 5835 | 467 | 4 | < 0.0001 | 0 | 0.7554 |
| NSP6 | 870 | 1162 | 3 | < 0.0001 | -3 | < 0.0001 |
| NSP12 | 2769 | 356 | 1 | 0.1789 | 2 | < 0.0001 |
| NSP14 | 1581 | 1781 | 1 | 0.8104 | 2 | 0.0129 |
| N | 1260 | 683 | 1 | 0.4540 | -1 | 0.0123 |
| NS7a | 366 | 223 | 0 | 0.5782 | 0 | 0.0458 |
| NSP13 | 1803 | 1659 | 0 | < 0.0001 | 0 | < 0.0001 |
| NS9b | 294 | 444 | 0 | 0.9704 | -4 | 0.0040 |
| E | 228 | 2338 | -2 | 0.1030 | -2 | 0.0486 |
| NSP2 | 1914 | 455 | -5 | 0.3511 | 3 | 0.0015 |
| NS9c | 222 | 2474 | -5 | 0.1000 | -7 | 0.0166 |
| NS3 | 828 | 243 | -5 | < 0.0001 | 0 | 0.0002 |
| NSP1 | 540 | 498 | -7 | < 0.0001 | -4 | 0.0001 |
| 3P | 370 | 188 | -9 | < 0.0001 | 0 | 0.9165 |
| RNA | 29903 | 744 | 27 | < 0.0001 | 3 | < 0.0001 |

**Figure 1:**
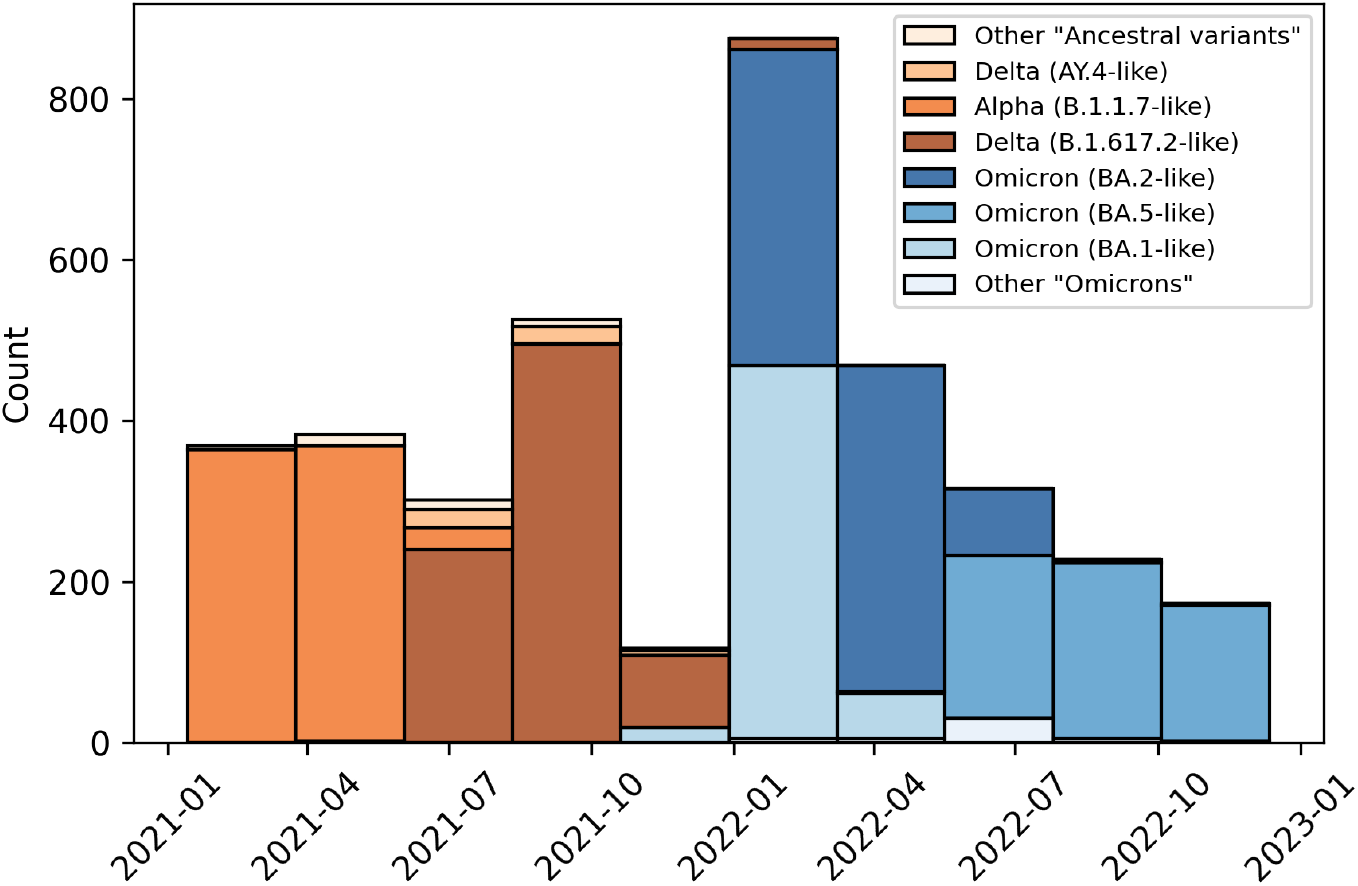
SARS-CoV-2 variants obtained from GISAID portal according to Berlin, Germany geotag.

### 2.2 Highly expressed miRNAs

miRNA-seq read count tables were downloaded from the GDC portal at https://portal.gdc.cancer.gov/projects/ for colon (COAD) and lung (LUAD) tissues. Following the corresponding clinical description, the dataset samples were referred to as “Normal” (“Solid Tissue Normal”) and “Tumor” (“Primary Tumor”) groups. Only “Normal” samples were used to assess miRNA expression in the analysis: 8 entries in COAD dataset and 46 in the LUAD project.

Highly expressed miRNAs were selected independently for each tissue by the following procedure: we summed up read counts associated with a particular miRNA by the samples; sorted miRNAs by the total number of reads; selected miRNAs, which account for 95% of all reads. As a final expression of the miRNAs, we averaged corresponding CPM expressions by the samples.

### 2.3 Binding model

To model the binding interaction between miRNA and viral mRNAs, the regions of viral RNA that were a reverse complement to the seed region of miRNA (nucleotides from 2 to 7 on the 5’-end of mature miRNA) were mapped to SARS-CoV-2 genomes. Following the common classification [20], such interactions are called “6mer”; they lead to translational inactivation of target mRNAs [21]. In addition to the reverse complementarity of the seed region, miRNA target prediction tools may rely on additional features. For example, TargetScan [22] utilizes context++ score based on site type, 3’-supplementary pairing, local AU content and others, MirTarget [23] considers GC content of target site, and UTR length, while RNA22 [24] predicts target sites by placing “Target Islands” within 3’-UTRs, 5’-UTRs or CDS.

Commonly, the overlap of the targets by different software is weak at best [25, 26]. While RNA22 predicts more individual miRNA-mRNA interactions than TargetScan, it misses the majority of classical seed region binding sites in 3’-UTRs. Moreover, the prediction results heavily depend on the tool-specific internal thresholds, with sensitivity and specificity expected to vary depending on particular thresholds. To overcome these limitations, we utilized a bottom-up approach based on the mapping of SARS-CoV-2 regions with reverse complement “6mers” matching the seed regions within tissue-specific and relevant host miRNAs.

All SARS-CoV-2 RNA sequences were aligned to the reference “Wuhan” variant using MAFFT tool [27]. After the alignment, we transformed viral RNAs coordinates on the “Wuhan” variant and annotated protein-coding regions using “Wuhan” annotation (https://www.ncbi.nlm.nih.gov/nuccore/1798174254).

### 2.4 Statistical analysis

In all comparisons, the significances of the findings were evaluated in the nonparametric Mann–Whitney U tests [28]. In addition, the standard Student’s t-test procedure [29] was used to test the hypothesis that the Spearman correlation [30] was zero.

The Monte-Carlo sampling procedure was used to show that the decrease in the occurrence of miRNA binding motifs in the“Omicron” sequence group was specific to the highly expressed miRNAs. More precisely, we fixed the expression profile of the highly expressed miRNAs and randomly replaced the miRNA sequences with other miRNAs expressed in a tissue. After that, we calculated the weighted number of binding motifs in “Omicrons” and “Ancestral” groups: the number of binding motifs within a particular miRNA was multiplied by the proportion of its read counts, summed up and normalized by the total lengths of the compared viral genomes. Then, for “Ancestral variants” and “Omicrons” groups, the computed numbers were compared using the Mann-Whitney U test. The described procedure was repeated 1,000 times. Thus, we obtained the distribution of statistics characterizing the change in the number of binding motifs between “Ancestral variants” and “Omicrons”. Finally, using this distribution, we calculated the probability that the decrease in the number of binding motifs for highly expressed miRNAs was greater than for randomly selected ones.

### 2.5 Other technical details

The statistical analysis (hypothesis testing, correlation computation) was performed with SciPy library [31]. All vector computations were implemented using NumPy library [32] and parallelized using Parallel tool [33]. Miscellaneous computations with tables were produced with Pandas [34]. Finally, figures were made using Matplotlib [35] and Seaborn [36] libraries. The source code of the analysis can be found at https://github.com/zhiyanov/covid-miRNA-evolution.

## 3 Results

In this study, both the tissue-specificity and expression volume restrictions were applied to facilitate identification of the most prominent interactions between host miRNAs and the viral genomes. First, the scope of the study was limited to the lungs and colon tissues as the two main sites of SARS-CoV-2 replication in the human body [15]. Second, we aimed to analyzed only those miRNA species that represent the top 95% of the miRNAome for their tissue type [37], with the following rationale. In COVID-19, each infected cell expresses from 1,500 to 15,000 copies of the virus [38] and from 100,000 to 200,000 miRNA molecules [39]. To visualize the kinetics of their interactions, one should take into account that each miRNA species, in addition to the virus, binds to 25-50 host mRNAs species, each expressed as 50 to 100 copies per cell, thus, and therefore, being supplied with more than 1,250 endogenous target molecules [21]. Because of that, only highly represented miRNA species are likely to exert a significant impact on the evolution of the viral genomes.

For lung tissue, there were 32 miRNAs (Figure 2 of Supplementary materials), and for the intestines, 40 miRNAs (Figure 3) that fulfilled the abovementioned criteria. Twenty-one of these miRNAs were found in both of these tissues, and 19 were expressed in either one or another tissue. In lung tissue, lung-specific miRNAs accounted for 81% of all miRNAs, while in the colon only 56% of expressed miRNAs were colon-specific. In lungs, the most prevalent miRNA was miR-143-3p, followed by miR-21-5p, miR-22-3p and miR-30a-5p. In colon, miRNA expression profile had a predominance of let-7b-5p, miR-92a-3p and miR-200c-3p.

**Figure 2:**
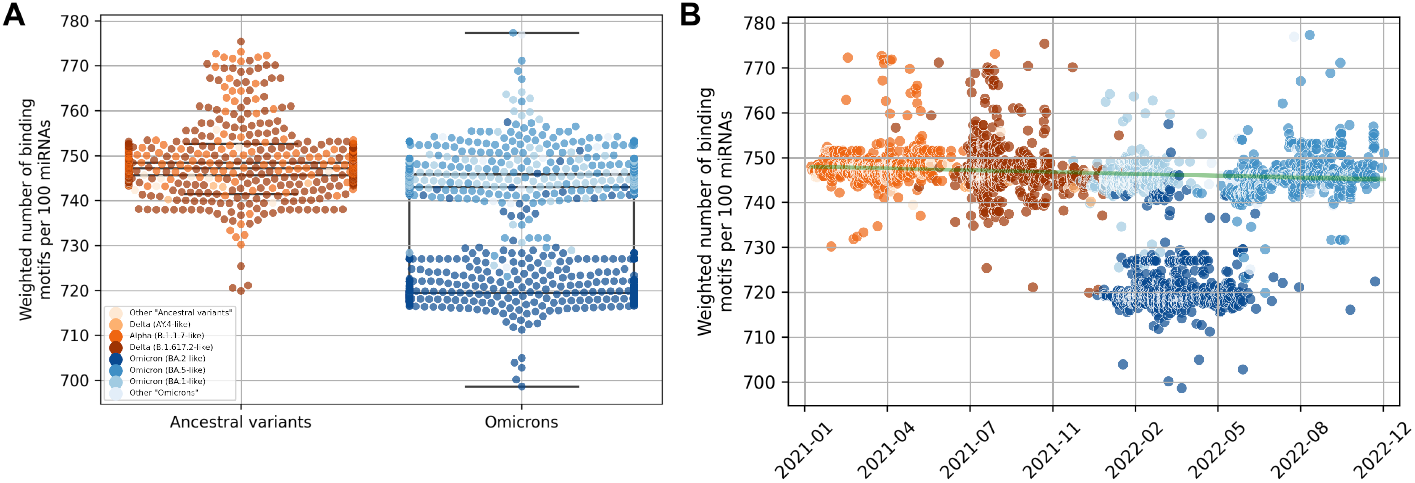
Weighted number (by miRNA expression in colon tissues) of binding motifs per 100 miRNA and 29,903 nucleotides **A**: grouped by “Ancestral variants” and “Omicrons”, **B**: distributed over time. Spearman correlation between the weighted number and number of days since the start of pandemics is ≤ −0.34, (*p* ≤ 7.37 · 10^-80^). In this analysis, Omicron (BA.2like) variants were excluded.

**Figure 3:**
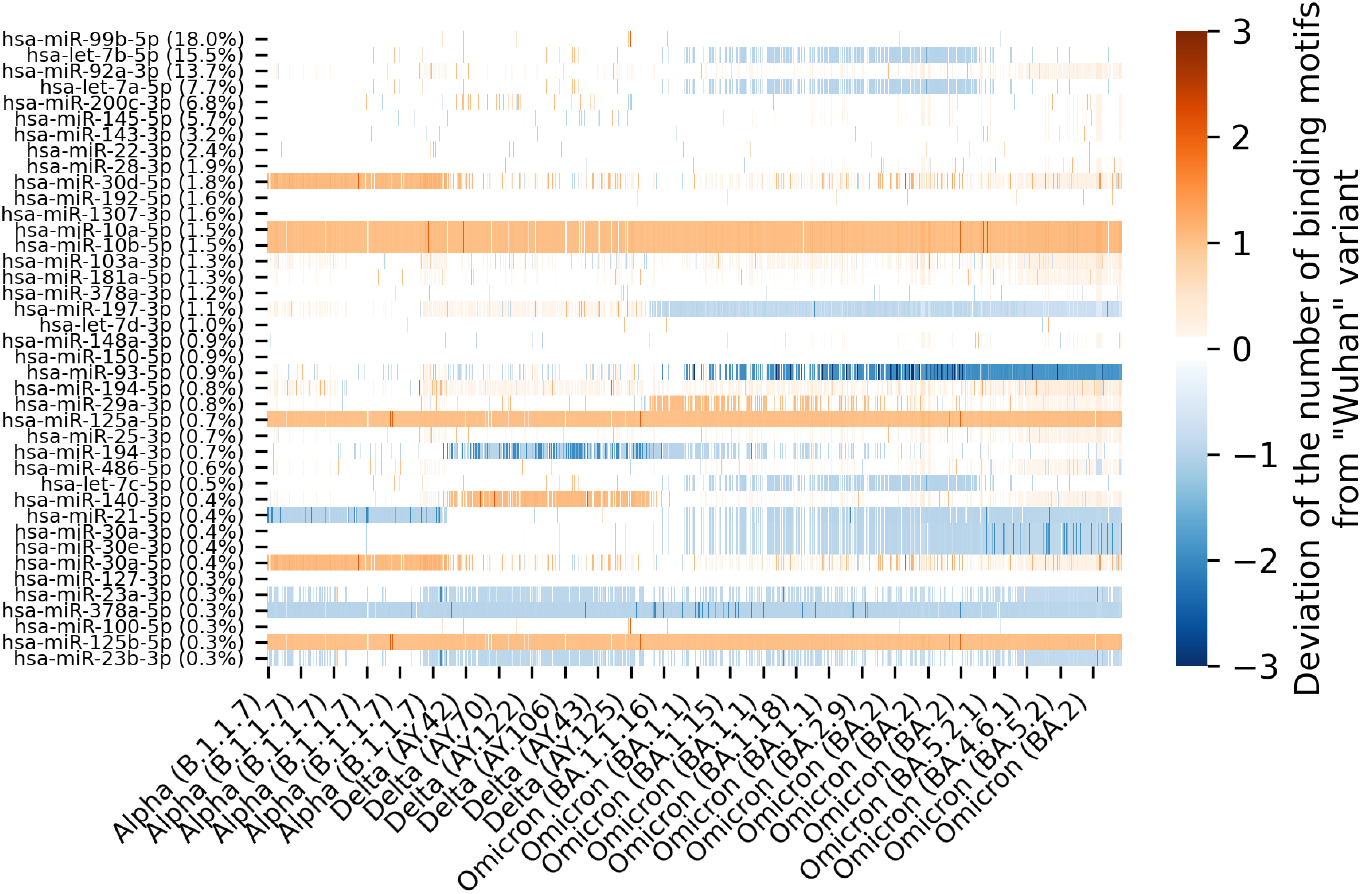
Amounts of the binding motifs in analyzed SARS-CoV-2 variants and reference “Wuhan” variant differ. Here, miRNAs are sorted by the levels of their expression in the intestine. The share of miRNome represented by particular miRNA could be found adjacent to miRNA ID.

From the GISAD database, a total of 3,755 SARS-CoV-2 sequences were retrieved. All of them were tagged by location within the same metropolitan area (Berlin, Germany) and a time stamp between the 1^st^ January 2020 and the 20^th^ December 2022 (Figure 1), see “On choosing a city of the study” section of Supplementary materials for more details). All selected sequences were divided into two groups:

- “Ancestral variants”, which included “Alpha” (B.1.1.7-like, 759 variants), “Beta” (B.1.351-like, 4 variants), Delta (B.1.617.2-like, 841 variants; AY.4.2-like, 12 variants) as well as 7 other strains (Table 1 of Supplementary materials). All of these sequences were collected in January 2022 and earlier; “Ancestral variants” included “Alpha” (B.1.1.7-like, 759 variants), “Beta” (B.1.351-like, 4 variants), Delta (B.1.617.2-like, 841 variants; AY.4.2-like, 12 variants) and 7 other strains (Table 1 of Supplementary materials) and belonged to the period until January 2022;
- “Omicrons”, which included 8 varieties of “Omicron” strain. All of these were collected in December 2021 or later.

To assess the evolutionary pressure exerted by colon and lung-specific miRNAs, we analyzed the expression-normalized amounts of binding motifs numbers within 29,903 nucleotides of viral SARS-CoV-2 RNA (see “Materials and methods” for more details). In this way, we took into account the regulatory contribution of each miRNA separately.

As compared to “Ancestral variants”, Omicron variants contain strikingly lesser amounts of binding sites for either colon miRNAs (Mann-Whitney U test, *p* ≤4.44 · 10^-254^, Figure 2, A) or lung miRNAs (Mann-Whitney U test, *p* ≤ 1.07 · 10^-146^, Figure 1 of Supplementary materials).

By reshuffling the list of miRNAs with a predefined expression repertoire specific to the highly expressed miRNAs, we calculated the proportion of random shuffles for which the drop in the number of binding motifs was greater than the drop detected for the highly expressed miRNAs (the detailed description of the procedure can be found in “Materials and Methods” section). We found that this proportion was lower than 5% for colon miRNAs (*p* ≤ 2.50 · 10^-2^), but not for lung miRNAs (*p* ≥ 7.79 · 10^-2^). When the significance of the miRNA influence was tested for other tissue compartments, including the prostate, the breast, the bladder, and the liver, the decrease in the number of observed motifs was not significant for either of these tissues (Table 2 of Supplementary materials).

Further study of the colon-specific dropdown showed that it was mainly observed within Omicron brunch (BA.2-like) (see Figure 2, A). However, even after exclusion of Omicron (BA.2-like) from “Omicrons” group, the indicated trend remained strong (Mann-Whitney U test, *p* ≤ 2.57 · 10^-69^). Moreover, the weighted number of miRNA binding motifs decreased steadily according to time stamps (Spearman correlation ≤ −0.34, *p* ≤ 7.37 · 10^-80^, Figure 2, B), thus, pointing at continuous evolution of “Omicron” sequences within the intestinal niche.

We calculated the difference between the number of binding motifs in the “Wuhan” and subsequent variant (Figure 3). It turned out that the loss of the binding sites was primarily due to diminished binding to hsa-let-7b-5p, hsa-let-7a-5p and hsa-miR-93-5p. Notably, for hsa-miR-194-5p and hsa-miR-29a-3p, the amounts of binding sites in the “Omicrons” have increased.

The pairwise alignment of SARS-CoV-2 RNAs with reference “Wuhan” variant allowed us to compare the distributions of the miRNA seed regions within each variant of the virus. Further, we calculated the binding distributions for individual miRNA, and found that the contribution of each miRNA in those distributions was proportional to its expression in colon tissues. Finally, the distributions were averaged over “Ancestral variants” and different groups of “Omicrons”, either including or excluding Omicron (BA.2-like) variants, and analyzed.

As seen from Figure 4, the decrease in amounts of the complementary seeds observed in Omicron BA.2-like variants was primarily due to a loss of hsa-let-7b-5p binding motifs located within NSP4. Indeed, C9861T nucleotide mutations occurred mostly in Omicron BA.2-like sequences (872 variants of 887), while the 9861^st^ position in all “Ancestral variants” sequences retained the cytosine. Interestingly, that transition was silent as it has not lead to amino acid mutation: both CTC and CTT triplets encode for lysine.

**Figure 4:**
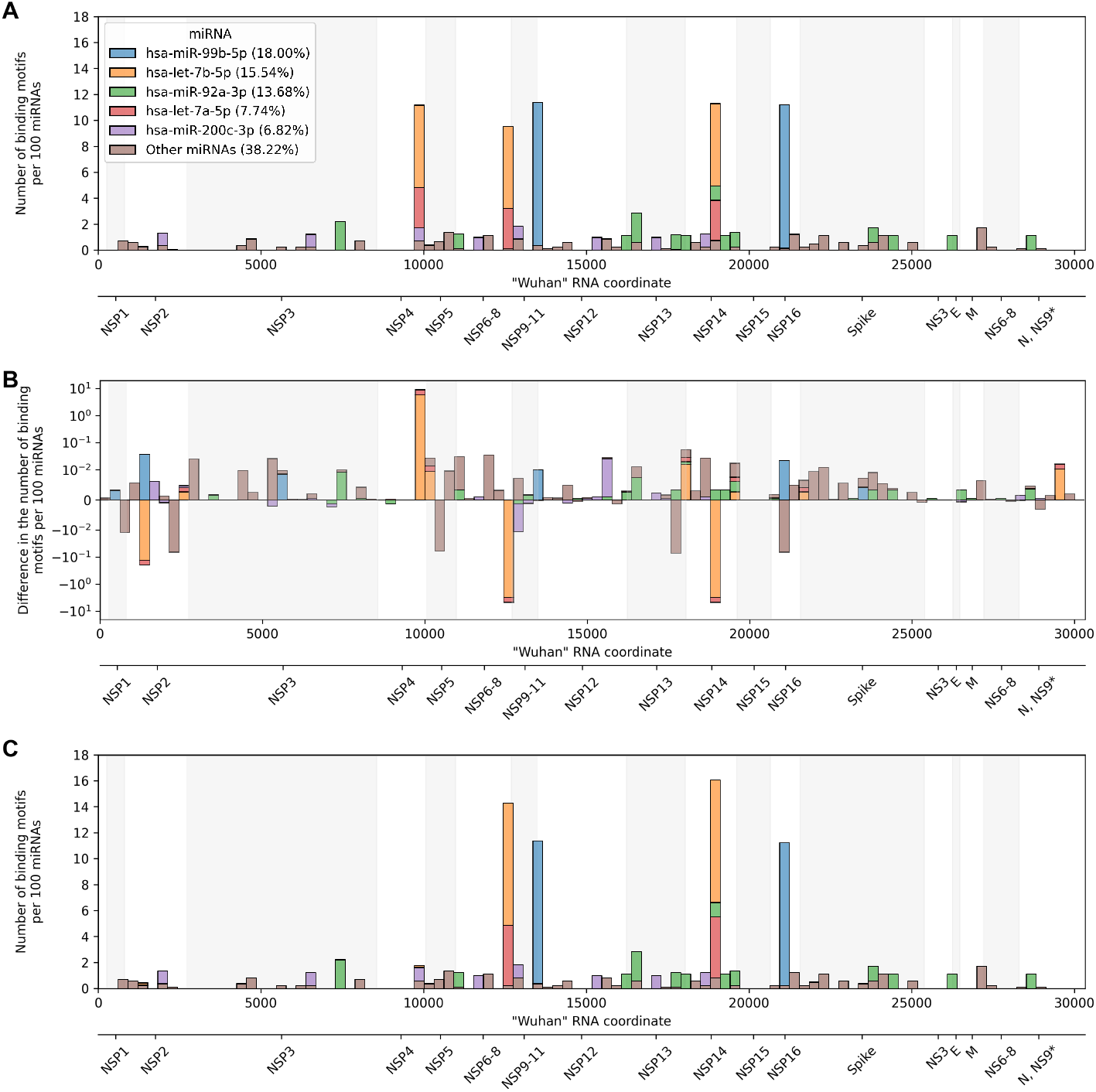
Distributions of miRNA binding motifs along to “Wuhan” RNA sequences and averaged by **A**: “Ancestral variants”, **B**: difference between “Ancestral variants” and Omicron (BA.2-like) variants, and **C**: Omicron (BA.2-like) variants.

Notably, a comparison of “Ancestral variants” and other “Omicrons” (Figure 3 of Supplementary materials) showed that the overall dropdown of miRNA binding sites was not due to any specific mutation within a particular miRNA binding motif. On the contrary, we observed a gradual decrease in the number of binding motifs (Figure 2, B) over time.

To access the contribution of virus RNA coding regions to the observed dropdown, the weighted (by miRNA expression) amounts of miRNA binding motifs per coding region were calculated for “Ancestral variants” and for “Omicrons” with or without Omicron (BA.2-like) sequences separately. By comparing these amounts within “Ancestral variants” with “Omicrons” using Mann-Whitney U test (Table 1), we found out that Spike and NSP15 regions were changed the most. In addition, the sequences classified as Omicron (BA.2-like) displayed an additional region with accumulated changes, the one encoding for NSP4, while in other “Omicrons” the third most divergent location was the region of NSP12.

## 4 Discussion

In this study, we tested the hypothesis that host cell miRNAs may have influenced the evolution of SARS-CoV-2. Here we studied two sets of miRNAs expressed in the main sites of virus reproduction, the lungs and the intestine, and found that these sets significantly differ in the representation of individual miRNAs. On one hand, hsa-miR-143-3p, hsa-miR-21-5p, hsa-miR-22-3p and hsa-miR-30a-5p were specific for the lungs, with large predominance of miR-143-3p expression (41.5%). On the other hand, the intestine was characterized by a more uniform profile with the presence of tissue-specific let-7b-5p, miR-92a-3p and miR-200c-3p.

We show that in course of pandemics, SARS-CoV-2 genomes lost many miRNA binding sites specific for the lung and intestinal tissue expressed miRNAs.

Importantly, Monte-Carlo simulations show that observed changes were significant only for the set of miRNAs expressed in the intestinal tissue (*p* ≤ 2.50 · 10^-2^).

Coronaviruses are generally perceived as Acute Respiratory Infections (ARIs), and the viral production in the lungs provides for a reservoir underlining the virus spread. In addition to the lungs, SARS-CoV-2 is capable of replication in many other organs and tissues [40], the main of which is an intestine, where the virus may reside for up to three month [41]. Prolonged period of intestinal replication provides sufficient time for evolutionary forces to work on the population of the viruses within the same host, with the host factors serving as the drivers for such evolution. Arguably, binding of host miRNAs represents such a force. Resultant viral populations may be returned to circulating pool of the viruses either through the fecal-oral route [42] or during secondary viremia in the immunocompromised hosts capable of supporting internal reinfection of the lungs [43, 44], and subsequent airborne spread.

It has been shown that the initial version of Omicron variant shared a high similarity with the variants of the SARS-CoV-2 virus found in rodents, particularly in mice [45]. Therefore, it has been hypothesized that one of the ancestral variants of SARS-CoV-2, from approximately mid-2020 [46], has entered the population of rodents and has evolved within these animals for quite some time till reemerging in humans [47, 48]. An alternative theory suggests a long-term virus evolution within an immunocompromised patient, predominantly in the intestines [49].

An observation described in this study highlights the significance of the intestinal site for the pathogenesis of COVID-19. Although pneumonia and respiratory complications account for the most of the morbidity and mortality of COVID-19, extrapulmonary manifestations of COVID-19 infection include prominently diarrhea, as well as nausea, emesis, anorexia, abdominal pain, and heartburn [41]. These complications are observed in 10%-20% of patients with COVID-19 [50], which does not exclude the possibility of asymptomatic gastrointestinal infections with SARS-CoV-2. Moreover, the latent persistence of SARS-CoV-2 in the intestine may be one of the contributors to the long COVID [51].

A dramatic decrease in miRNA binding sites was especially pronounced in BA.2-like SARS-CoV-2 variants. To a large extent, this drop was due to the mutations within the region encoding the NSP4 protein, which promotes the formation of double-membrane vesicles serving as a scaffold for replication-transcriptional complexes [52] of the virus. Additionally, attachment to NSP4 protects viral RNA from intracellular immunity [53].

Along with dramatic changes in S-protein, the miRNA-driven alterations of NSP4, acquired during prolonged replication with the intestine, may have contributed to the current global dominance of the descendants of the BA.2 variant [54]. Our findings highlight the possibility that intestinal tissue may significantly impact evolution of the SARS-CoV-2 genome and may play a pivotal role in the long COVID.

## 5 Acknowledgements

The authors thank Dr. Stepan Nersisyan from Computational Medicine Center at Thomas Jefferson University for useful comments and discussions.

## 6 Conflict of interest

Alexander Tonevitsky is an employee of Art Photonics GmbH.

## 7 Funding

The research was performed within the framework of the “Creation of Experimental Laboratories in the Natural Sciences Program” and Basic Research Program at HSE University.

