## Supplementary materials for "miRNA binding pressure channels evolution of SARS-CoV-2 genomes"

### On choosing a city of the study

In this study, we analyzed the virus sequences within a particular city since tracking the change in virus variants within a whole country is almost impossible After the lifting of isolation, the movement of the population intensified New variants of the virus entered countries through major cities with airports and spread from there In addition to analyzing virus sequences within one locality, we made high demands on the quality of sequences: the absence of undefined nucleotides and a complete sequence length This is due to the specifics of the analysis methodology We analyzed 6-mers, which are extremely sensitive to substitutions and sequence length GISAID entries from London (UK), Berlin (Germany) and Taiwan were analyzed The highest number of qualitative sequences was in Berlin They were used in further analysis.

### Materials

| Strain | Initial | Selected |
|---|---|---|
| Alpha (B.1.1.7-like) | 3791 | 759 |
| B.1.1.318-like | 10 | 0 |
| B.1.1.7-like + E484K | 87 | 17 |
| B.1.617.1-like | 16 | 2 |
| Beta (B.1.351-like) | 102 | 4 |
| Delta (AY.4-like) | 736 | 49 |
| Delta (AY.4.2-like) | 73 | 12 |
| Delta (B.1.617.2-like) | 8625 | 841 |
| Delta (B.1.617.2-like) + K417N | 7 | 2 |
| Eta (B.1.525-like) | 155 | 2 |
| Gamma (P.1-like) | 24 | 1 |
| Lambda (C.37-like) | 8 | 2 |
| Mu (B.1.621-like) | 3 | 0 |
| Omicron (BA.1-like) | 2251 | 539 |
| Omicron (BA.2-like) | 5736 | 887 |
| Omicron (BA.4-like) | 204 | 28 |
| Omicron (BA.5-like) | 4285 | 591 |
| Omicron (Unassigned) | 767 | 9 |
| Probable Omicron (BA.1-like) | 86 | 2 |
| Probable Omicron (BA.2-like) | 19 | 3 |
| Probable Omicron (BA.4-like) | 1 | 0 |
| Probable Omicron (BA.5-like) | 13 | 5 |
| Probable Omicron (Unassigned) | 32 | 0 |

Table 1: SARS-CoV-2 strains distributed in Berlin of Germany during the period from 01.01.2020 to 20.12.2022. “Selected” column contains the number of high-quality RNA variants selected for the analsysis from “Initial” ones.

| Tissue | $p$-value | $p$-adj |
|---|---|---|
| Breast (TCGA-BRCA) | 0.070 | 0.280 |
| Prostate (TCGA-PRAD) | 0.117 | 0.234 |
| Bladder (TCGA-BLCA) | 0.166 | 0.221 |
| Liver (TCGA-LIHC) | 0.189 | 0.189 |

Table 2: Results of the Monte-Carlo sampling procedure for miRNA profiles of TCGA-project normal samples of different tissues.

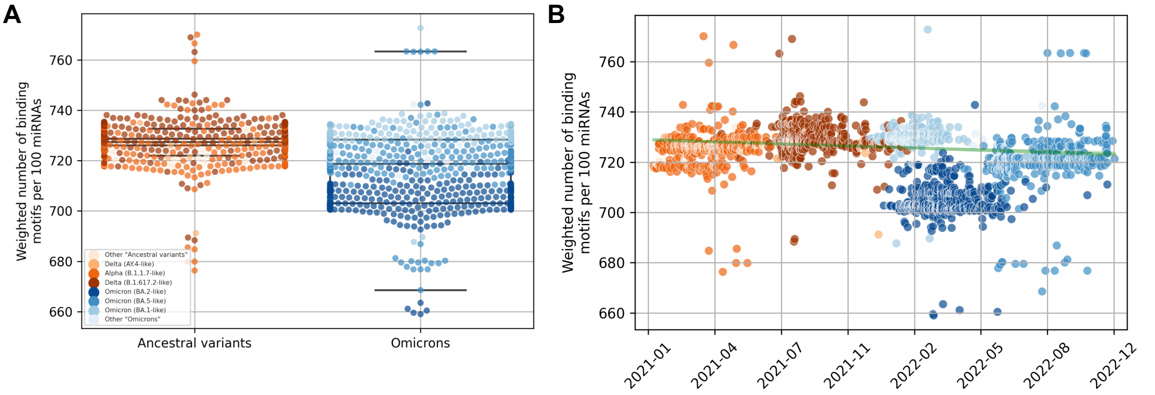


Figure 1: Weighted number (by miRNA expression in lung tissues) of binding motifs per 100 miRNA and 29,903 nucleotides **A**: grouped by "Ancestral variants" and "Omicrons", **B**: distributed over time. Spearman correlation between the weighted number and number of days since the pandemic start (Omicron (BA.2-like) variants are excluded) equals $-0.10$ ($p \leq 4.54 \cdot 10^{-8}$).

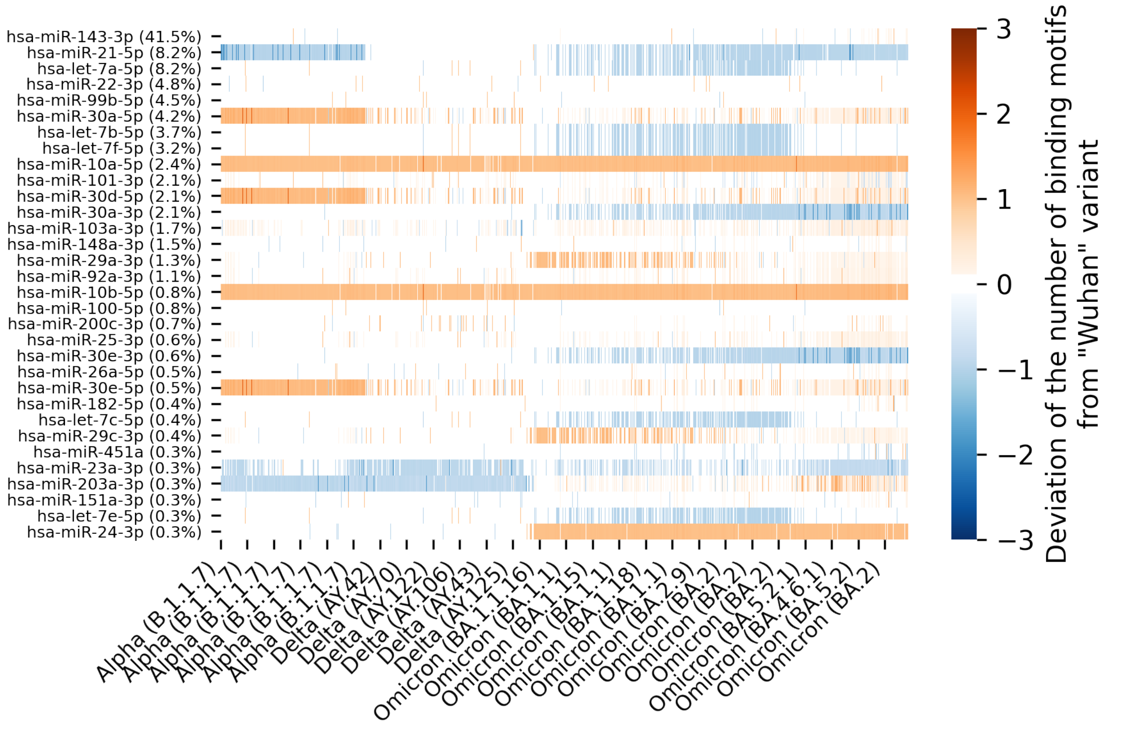


Figure 2: Difference of the binding motifs number between analyzed SARS-CoV-2 variants and reference "Wuhan" variant. MiRNAs are sorted by their expression in lung tissues. The percentage of miRNA expression is located next to its name.

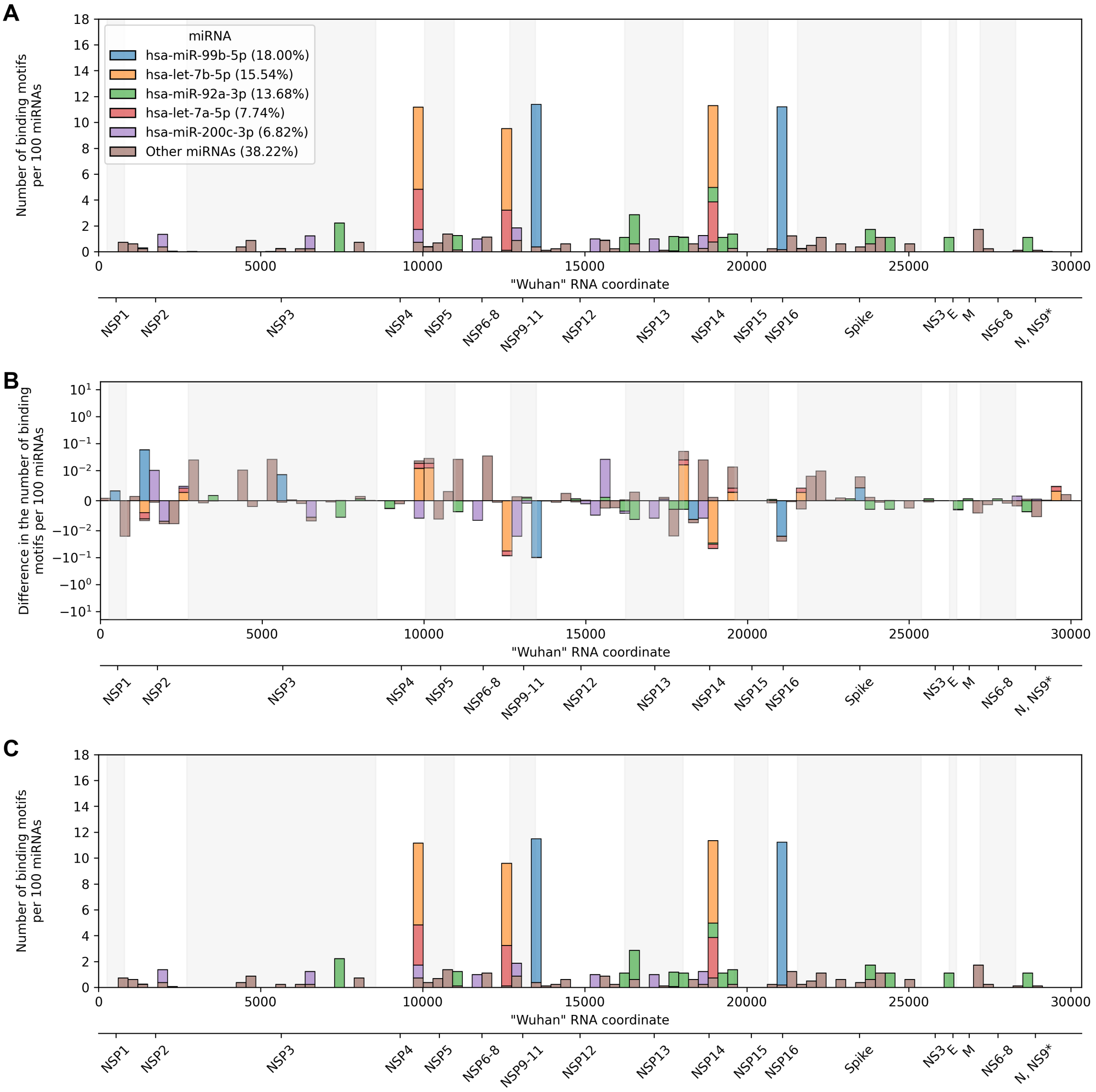


Figure 3: Distributions of colon miRNA binding motifs transformed to "Wuhan" RNA and averaged by **A**: "Ancestral variants", **B**: difference between "Ancestral variants" and "Omicrons" excluding Omicron (BA.2-like), and **C**: "Omicrons" excluding Omicron (BA.2-like).

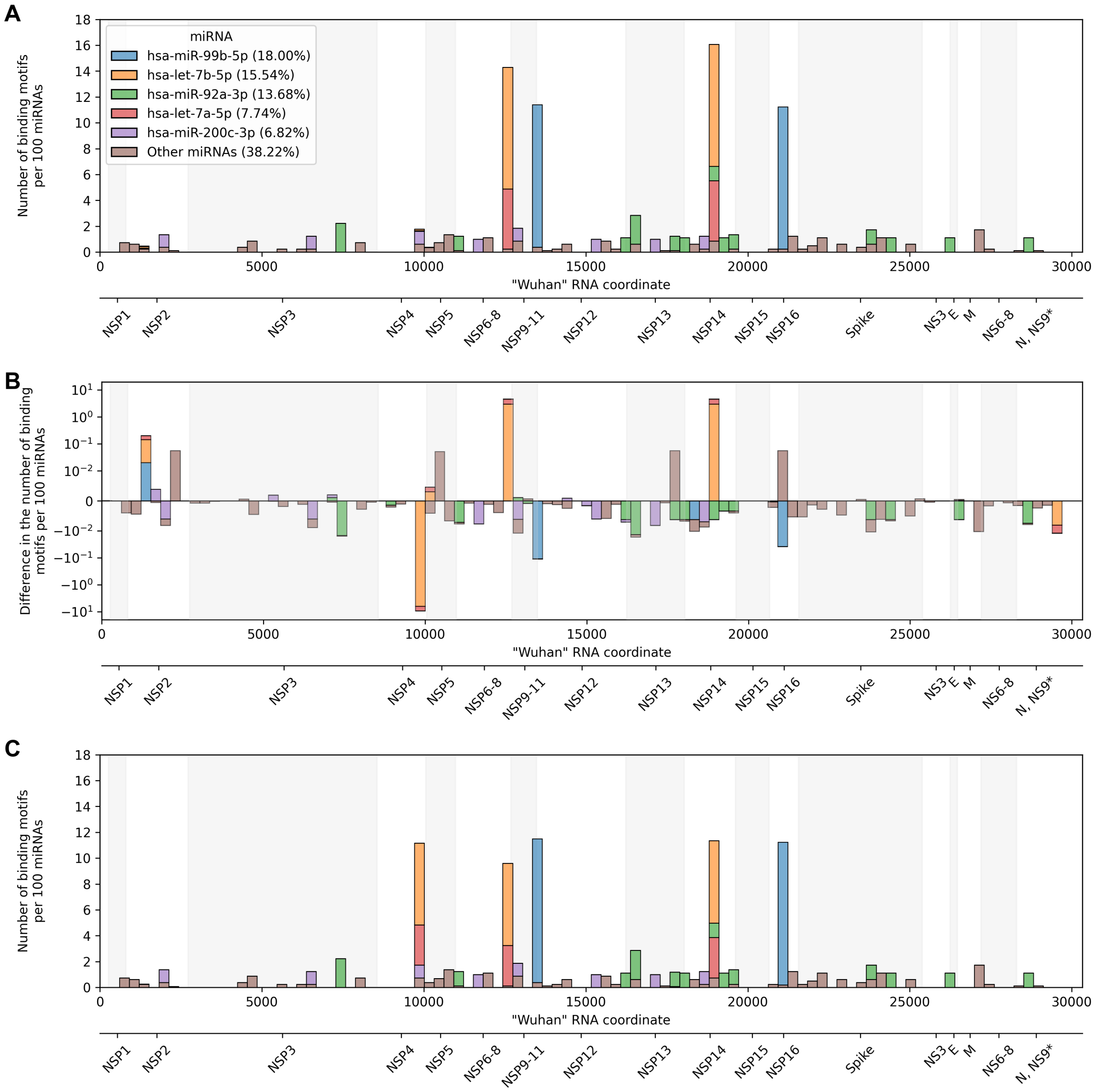


Figure 4: Distributions of colon miRNA binding motifs transformed to "Wuhan" RNA and averaged by **A**: Omicron (BA.2-like) variants, **B**: difference between Omicron (BA.2-like) variants and other "Omicrons", and **C**: other "Omicrons".
